# Oral Pretreatment with Galantamine Effectively Mitigates the Acute Toxicity of a Supra-Lethal Dose of Soman in Cynomolgus Monkeys Post-treated with Conventional Antidotes

**DOI:** 10.1101/2020.02.03.932798

**Authors:** Malcolm Lane, D’Arice Carter, Joseph D. Pescrille, Yasco Aracava, William P. Fawcett, G.W. Basinger, Edna F.R. Pereira, Edson X. Albuquerque

## Abstract

The present study was designed to evaluate the effectiveness of galantamine administered orally as a pre-treatment to mitigate the acute toxicity of 4.0xLD50 soman in Cynomolgus monkeys post-treated with atropine, 2-PAM, and midazolam. Pharmacokinetic experiments revealed that the oral doses of 1.5 and 3.0 mg/kg galantamine HBr were quickly absorbed and produced plasma concentrations of galantamine that generated approximately 20% to 40% reversible inhibition of blood acetylcholinesterase (AChE) activity. This degree of reversible AChE inhibition has been shown to be safe and sufficient to protect AChE from the irreversible inhibition by nerve agents, and, thereby, suppress the acute toxicity of these agents. Thus, in subsequent experiments, adult male Cynomolgus monkeys were pretreated orally with 1.5 or 3.0 mg/kg galantamine, challenged intramuscularly with 4.0xLD50, and post-treated with intramuscular injections of 0.4 mg/kg atropine, 30 mg/kg 2-PAM, and 0.32 mg/kg midazolam. All animals subjected to these treatments survived the soman. By contrast, none of the animals that were pretreated with saline and only 40% of the animals that were pretreated with pyridostigmine survived the soman challenge when post-treated with the same conventional antidotal therapy as that delivered to the galantamine-pretreated, soman-challenged monkeys. In addition, large numbers of degenerating neurons were visualized in the hippocampi of soman-challenged monkeys that had been pretreated with pyridostigmine or saline, but not in the hippocampi of animals that had been pretreated with galantamine. To our knowledge, this is the first study to demonstrate the effectiveness of clinically relevant oral doses of galantamine to prevent the acute toxicity of supra-lethal doses of soman in non-human primates.

## Introduction

Nerve agents, including soman, sarin, VX, and novichock, are highly toxic organophosphorus (OP) compounds that have been used during the Second Sino-Japanese War in 1937-1945, the 1980s Iran-Iraq conflict, the 1995 Tokyo subway terrorist attack, the 2013 Syrian Civil War, and the recent 2017-2018 terrorist attacks against civilians in Syria, Malaysia, and the United Kingdom (rev. in Romano and King, 2001; Franca et al., 2019). The acute toxicity triggered by nerve agents and their chemically related OP insecticides can be fatal and results primarily from overstimulation of muscarinic and nicotinic receptors by acetylcholine (ACh) that accumulates in the peripheral and central nervous systems as OP compounds irreversibly inhibit acetylcholinesterase (AChE), the enzyme that catalyzes ACh hydrolysis. Thus, subjects poisoned with OP compounds develop a cholinergic crisis typically characterized by miosis, profuse secretions, diarrhea, bronchoconstriction, muscle fasciculation, tremors, and seizures (reviewed in Hurst et al., 2012).

The standard therapy used to treat OP intoxication includes atropine to block overactivation of muscarinic receptors, an oxime (generally 2-PAM) to reactivate AChE, and a benzodiazepine (midazolam or diazepam), as needed, to suppress seizures (reviewed in Newmark, 2019). However, this conventional antidotal therapy has some important limitations. Specifically, AChE inhibited by some OP compounds, particularly soman, become quickly refractory to reactivation by oximes (Marrs et al., 2006), and benzodiazepines progressively lose their anticonvulsant effectiveness if treatment is delayed after onset of OP-induced seizures (McDonough et al., 2010). Consequently, concerted efforts are underway for development of more effective post-treatments (Jett and Laney, 2019). Equally important, however, is the need of an effective pretreatment to protect military personnel and first responders at risk of exposure to high levels of OP compounds in a chemically contaminated environment. During the 1995 Tokyo sarin attacks, numerous first responders were hospitalized with acute signs of toxicity due to their primary or secondary exposure to the nerve agent while attending the mass casualties (Nishiwaki et al., 2001). Yet, to date, personal protective equipment remain the only protective measure used by first responders attending OP poisoning victims (Chai et al., 2017).

In 2006, galantamine, a centrally acting reversible AChE inhibitor approved for treatment of mild-to-moderate Alzheimer’s disease, emerged as a potential effective and safe pretreatment against OP poisoning. Used as a stand-alone pretreatment, intramuscularly delivered galantamine prevented the acute toxicity of a lethal dose of nerve agents in guinea pigs and non-human primates (Albuquerque et al., 2006; Golime et al., 2018; Hamilton et al., 2017; Hilmas et al., 2009). Used as an adjuvant pretreatment and/or immediate post-treatment, intramuscularly delivered galantamine also increased the effectiveness of a high dose of atropine to mitigate the acute toxicity of supra-lethal doses of the OP insecticide paraoxon and the nerve agents soman, sarin, and VX in rodents (Albuquerque et al., 2006; Golime et al., 2018; Hilmas et al., 2009). The effectiveness of galantamine as a pretreatment against OP intoxication has been attributed, at least in part, to its ability to protect AChE in the peripheral and central nervous systems from OP-induced irreversible inhibition and to its neuroprotective properties (Pereira et al., 2010).

To date, no study has focused on the more practical use of an oral galantamine pretreatment to prevent the acute toxicity of nerve agents. The present study was, therefore, designed to evaluate whether an oral galantamine pretreatment effectively prevents the acute toxicity of a supra-lethal dose of soman in Cynomolgus monkeys post-treated intramuscularly with human-equivalent doses of atropine, 2-PAM, and midazolam that are recommended for use in pre-hospital settings. Results presented here reveal that this treatment regimen is more effective against the acute toxicity of soman than post-treatment with the conventional antidotes alone or in association with an oral pretreatment with pyridostigmine, a drug currently approved by the U.S. Food and Drug Administration (FDA) for use as pretreatment by military personnel at risk of exposure to soman (Aebersold, 2012). Based on the pharmacokinetic profile of the oral doses of galantamine studied here and the maximal degree of blood AChE inhibition they produced, the doses that protected Cynomolgus monkeys against soman poisoning translate into human equivalent doses known to be safe.

## Materials and Methods

### Subjects

Twenty 4-6-year-old male Cynomolgus monkeys weighing 3.69-6.89 kg were obtained from a commercial vendor (AlphaGenesis, Yemassee, SC, USA). The subjects were quarantined under the guidelines of the Centers for Disease Control for 3 months, during which time they were pre-screened to exclude potential tuberculosis and simian retrovirus infections. Animals were individually housed in stainless steel cages with water available ad libitum and visual and auditory contact with each other. The colony was maintained at 20-22°C with a relative humidity of 50% (±15%) on a 12-h light/dark cycle. Experiments were performed during the light phase. All experimental procedures were approved by the University of Maryland School of Medicine Animal Care and Use Committee and were conducted in full accordance with the National Institutes of Health (NIH) Guide for the Care and Use of Laboratory Animals.

### Pharmaceutical and Non-Pharmaceutical Chemicals

A stock solution of approximately 2.0 mg/ml soman (methylphosphonofluoridic acid 1,2,2-trimethylpropyl ester) was obtained through a Chemical Agent Provisioning Agreement between the University of Maryland, Baltimore and the U.S. Army Materiel Command (AMC). The stock solution was stored, handled, and disposed of according to standard operating procedures and following the regulations set forth by the U.S. AMC. Galantamine HBr and pyridostigmine bromide were purchased from Sanochemia Pharmazeutika (Vienna, Austria) and Sigma-Aldrich (St. Louis, MO, USA), respectively. Pharmaceutical grade solutions of saline (0.9% NaCl, injectable), atropine sulfate (15 mg/ml, injectable), and midazolam hydrochloride (5 mg/ml, injectable) were purchased from VetOne/MWI Veterinary Supply Co. (Boise, ID, USA), Sparhawk Laboratories, Inc. (Shawnee Mission, KS, USA), and Hospira, Inc. (Lake Forest, IL, USA), respectively. Pharmaceutical grade 2-PAM chloride for injection (Protopam®, lyophilized 1 g) was purchased from Baxter Healthcare Corp. (Deerfield, IL, USA). An injectable solution of ketamine (Zetamine®, 100 mg/ml ketamine HCl) was also purchased from VetOne/MWI Veterinary Supply Co.

### Oral Treatments with Test and Control Articles

Daily doses of 8 to 24 mg galantamine are currently approved for treatment of Alzheimer’s disease. Using the allometric scaling factor derived from body surface area of humans and monkeys (Reagan-Shaw et al., 2008), these human doses translate to approximately 0.41 mg/kg and 1.25 mg/kg galantamine base, respectively, or 0.52 mg/kg and 1.6 mg/kg galantamine HBr, respectively, in monkeys. Thus, the oral doses of galantamine studied here were 0.5, 1.5, and 3.0 mg/kg galantamine HBr, a range that includes and exceeds the human equivalent doses of this drug.

The oral dose of pyridostigmine studied here (i.e., 1.2 mg/kg) is the same as that which generated ∼40% reversible inhibition of red blood cell AChE in Rhesus monkeys (Hayward et al., 1990). This degree of enzyme inhibition has been shown to significantly increase survival of primates that are subsequently exposed to soman (see Dunn and Sidell, 1990 and references therein) and has, therefore, been used as the benchmark to establish the oral dose of pyridostigmine bromide to be used by military personnel at risk of exposure to soman (Dunn and Sidell, 1990).

Solutions of galantamine HBr (20 mg/ml) and pyridostigmine bromide (20 mg/ml) were prepared by quantitatively dissolving the salts in an appropriate volume of a pharmaceutical-grade solution of saline. The concentrations of both solutions were verified using the compendial methods described in the US Pharmacopeia. The volume of the solution needed to deliver a given dose of a test article (galantamine HBr or pyridostigmine bromide) or the control article (saline) was determined based on the animal’s body weight on the day of the experiment. A plain oatmeal cookie (Homekist™) divided into 4 quarters that were pre-impregnated with the adequate volume of a test solution was used to deliver the oral treatment to the animals in the morning following an overnight fast. Each animal was closely monitored to ascertain that it consumed the entire cookie. In case cookie crumbs were wasted by a monkey during treatment, they were collected on a mesh screen and weighed so that any wasted volume could be estimated and replaced with a new piece of cookie. Upon eating the required treat, animals received the remainder of their normal diet.

### Pharmacokinetic Experiment: Treatment and Blood Draws

Animals were randomly assigned to the different treatment groups, which consisted of a single oral dose of 0.5 mg/kg, 1.5 mg/kg, or 3.0 mg/kg galantamine HBr (n = 5, 10, and 5, respectively). Twenty-four h prior to a treatment, each animal was transferred to a stainless-steel squeeze-back cage (61 cm W × 71 cm D × 86 cm H) and transported to the treatment room, which was distant from the homeroom. After restraint, animals were sedated with 5-10 mg/kg ketamine (i.m.) and subsequently laid down on a procedure table. Their body temperature and respiratory rate were continuously monitored during sedation. One of the thighs was shaved and 1 ml of blood was collected by venipuncture from the saphenous vein into heparinized tubes. Following the blood draw, animals were transferred to the squeeze-back cage, where they were allowed to recover prior to being transferred to their home cage.

On the next day, animals were again transported to the treatment room, where they were treated, as described in the previous section, with an oral dose of galantamine HBr. Ten minutes after their oral treatment with galantamine, animals were sedated with ketamine (5-10 mg/kg, i.m.) and maintained under sedation, with a repeated ketamine dose, for the first hour of the pharmacokinetic experiment, enabling blood to be sampled by venipuncture at 20 min, 40 min, and 60 min after the treatment. Animals were allowed to recover from sedation and were then resedated with the same ketamine dose at each subsequent time point of blood sampling (2, 5, and 24 h post-treatment). At the end of the pharmacokinetic study, Cynomolgus monkeys were housed for one month prior to the beginning of the pharmacodynamic experiment.

### Measurements of Galantamine Concentrations in Plasma of Treated Cynomolgus Monkeys and Pharmacokinetic Analysis

Blood drawn from all animals were separated into 0.5 ml aliquots transferred to 2 heparinized vials, one of which was centrifuged to separate plasma, which was, then, stored as 200-µl aliquots. Plasma from all animals treated with 0.5 and 3.0 mg/kg galantamine and from 5 animals randomly selected from the 10 animals treated with 1.5 mg/kg galantamine was used for determination of plasma concentrations of galantamine by liquid chromatography in tandem with mass spectrometry (LC-MS/MS) following the methods described by Steiner et al., (2012) and Suresh et al. (2014). In short, 160 µl of plasma samples were spiked and thoroughly mixed with 40 µl of the internal standard solution containing 1.25 µg/ml diphenhydramine HCl (purity ≥ 99% by HPLC, Sigma-Aldrich, MO, USA). Following the addition of 1.4 ml of trichloromethane, the mixture was vortexed for 2 min and subsequently centrifuged at 2,000g. The organic phase was, then, evaporated under nitrogen at 45°C, and the remaining residue was reconstituted with 100 µl of the LC mobile phase consisting of 0.1% formic acid, 9.9% HPLC grade water, 90% methanol. Then, 20 μl of the reconstituted material were used for an LC-MS/MS analysis performed on Waters TQ-XS triple quadrupole mass spectrometer with ACQUITY UPLC platform (Thermo Scientific, San Jose, CA). The LC separation was performed on an ACQUITY-Xselect CSH C18 130Å (4.6 × 100 mm, 1.7 μm) (Waters Corp., Milford, MA) connected to an ACQUITY CSH C18 130Å VanGuard pre-column (2.1 x 5 mm; 1.7 µm) (Waters Corp., Milford, MA). Mobile phase A consisted of 0.1% formic acid in UPLC-grade water, and mobile phase B consisted of 0.1% formic acid in acetonitrile. The elution gradient program was as follows: 0-5 min, 2% → 33% of B; 5.1-7.0 min, 100% of B; 7.1-10.5 min, 100% of B → 2% of B, and the flow rate was set at 0.6 ml/min. Detection was performed in the positive-ion mode and positive electrospray ionization data were acquired using multiple reaction monitoring of the precursor to product ion (M+H)+ transitions of 288-to-213 m/z and 256-to-167 m/z for galantamine and diphenhydramine.

The area under the plasma concentration-time curve from time 0 to infinity (AUC0-∞) and half-life (t1/2) of galantamine were estimated through a non-compartmental analysis of the data using the linear-log trapezoidal fit in the software Kinetica 5.0 (Thermo Scientific Kinetica, Philadelphia, PA).

### Analytical Method for Determination of AChE Activity

Acetylcholinesterase (AChE) activity was determined in whole blood using a modification of the radiometric assay described by Johnson and Russell (1975). Briefly, 90 µl of whole blood samples pretreated with the butyrylcholinesterase inhibitor tetraisopropyl pyrophosphoramide (iso-OMPA, 0.1 mM final concentration) were incubated at room temperature for 2 min with 10 μl of 0.1 M ACh chloride and [acetyl-3H]ACh iodide (20 μCi/ml, which produced approximately 100,000 cpm when totally hydrolyzed by eel AChE). At the end of the incubation time, the reaction was stopped by the addition of an aqueous solution (100 μl) consisting of chloroacetic acid (0.50 M), sodium chloride (1 M), and sodium hydroxide (0.25 M). After vortexing the mixture, samples were clarified by centrifugation, and the clarified samples (140 μl) were transferred to a scintillation vial containing 3.86 ml of the fluor cocktail, which consisted of 90% (v/v) toluene, 10% (v/v) 3-metyl-1-butanol, 0.03% (w/v) 1,4-bis(5-phenyl-2-oxazolyl), and 0.05% (w/v) 2,5-diphenyloxazole. The mix was vortexed for 60 s, and the amount of tritiated acetate in the organic phase was measured by liquid scintillation counting for 2 min (Tri-Carb 2900TR, Perkin Elmer). Each sample was assayed in triplicate and counts were corrected for background by subtraction of counts obtained using whole blood samples devoid of cholinesterase activity and treated in a similar fashion. Background-corrected counts obtained from the blood sample drawn from each animal prior to its treatment with galantamine were taken as 100% and used to normalize the background-corrected counts obtained from blood samples drawn at different times after the treatment of that animal. Percentage of inhibition was then taken as 100% - normalized AChE activity at each time point.

### Pharmacodynamic Experiment: Treatments, Soman Challenge, Functional Observation Battery, and Survival Data Analysis

Animals were randomly assigned to one of four experimental oral pretreatments: saline (0.15 ml/kg; n = 4), 1.5 mg/kg galantamine (n = 5), 3.0 mg/kg galantamine (n = 6), or 1.2 mg/kg pyridostigmine (n = 5). On each experimental day, animals were transferred to a stainless-steel squeeze-back cage and transported to the treatment room, where they were orally treated, as described above, with a test or control article. One hour after the oral treatment, each animal was constrained with the help of the squeeze mechanism, and soman (4.0xLD50 or 15.08 µg/kg) was administered intramuscularly in the quadriceps muscle. Immediately after the nerve agent challenge, animals were post-treated intramuscularly with atropine sulfate (0.4 mg/kg), 2-PAM chloride (30 mg/kg), and midazolam (0.32 mg/kg) in rapid succession. The doses of the antidotes are equivalent, based on allometric conversion, to those recommended for treatment of OP poisoning in pre-hospital settings.

After the injections, the squeeze mechanism was partially released, and animals were closely monitored. A Functional Observation Battery (FOB) described earlier (Gauvin and Baird, 2008) was recorded at fixed time intervals before pre-treatment, between pre-treatment and the soman challenge, and after antidotal treatment. The FOB was recorded by two assistants, who worked independently, were unaware of the pre-treatment given to the animals, and were involved neither in animal restraint nor in the injection procedure. The FOB consisted of scoring the animal’s posture, arousal, vocalization, stereotypical behavior, facial movements, visual and auditory responses, gross motor movements, ocular changes (nystagmus, pupil dilation), leg muscle fasciculations, and eye lid paralysis, in addition to the typical clinical signs of a cholinergic crisis, including salivation, lacrimation, bronchial secretions, increased urination, diarrhea, tremors, convulsions, and breathing distress. Animals were euthanized humanly, as described below, at 24 h after the soman challenge or at the time they presented life-threatening signs of toxicity, which included unremitting convulsions that lasting longer than 20 min, loss of body temperature, loss of posture, and severe respiratory distress characterized by gasping. Prior to euthanasia, a veterinarian confirmed that an animal had reached the endpoint for euthanasia. Number of surviving animals in each group was plotted against time and compared among the treatment groups using a Kaplan-Meier analysis.

### Histopathological Analysis

At 24 h post-soman challenge or at the time life-threatening signs of toxicity were evident, animals were sedated with ketamine (10 mg/kg, i.m.), transferred to the procedure table, and given an intravenous injection of heparin to prevent blood coagulation. Following the heparin injection, animals were euthanized with an i.m. overdose of ketamine and transcardially perfused with phosphate buffered saline (PBS, pH = 7.4) followed by 4% paraformaldehyde in PBS. Their brains were, then, harvested, post-fixed overnight at 4°Cwith freshly prepared 4% paraformaldehyde in PBS, cryoprotected with a solution of 30% sucrose in PBS, and frozen with crushed dry ice before storage at −80°C.

Using a Leica CM 3000 cryostat (Leica Biosystems, Buffalo Grove, IL), frozen brains were cut in 30-µm thick slices, which were, then, transferred to wells containing PBS. This was followed by three washes in 70% ethanol, 50% ethanol, and water, each for 1 min. After being washed in PBS and water, slices were immersed in a potassium permanganate solution (0.06%) for 15 min, on a rotating platform, rinsed in water for 1 min, and finally transferred to wells containing the Fluoro Jade-B (FJB, Chemicon, Temecula, CA, 0.0004%) staining solution prepared as described in Mamczarz et al. (2016). Slices were incubated with FJB for 30 min, under gentle shaking in the dark.

After being rinsed with water, slices were transferred to a microscope slide, dried on a slide warmer (50°C), immersed in xylene, and cover slipped with Cytoseal® mounting media (Richard Allan Scientific, Kalamazoo, MI). Slides were imaged with a Nikon Eclipse 80i upright microscope equipped with a Ds-FiZ camera controlled by NIS-Elements BR 3.0 SP4 software (Nikon Instruments Inc., Melville, NY). The fluorescence intensity was visualized using a fluorescein isothiocyanate filter set (Excitation: 465 nm; Emission: 520 nm).

## Results

### Plasma Levels of Galantamine Following Oral Treatment of Cynomolgus Monkeys with Galantamine HBr

As shown in Figure 1, following the oral administration of 0.5, 1.5, or 3.0 mg/kg galantamine HBr to Cynomolgus monkeys, plasma levels of galantamine increased up to a maximum between 1 and 3 h post-treatment and declined subsequently. One of the five animals treated with 3.0 mg/kg galantamine had no measurable plasma concentrations of galantamine. Therefore, data from this animal were not included in the analysis.

**Figure 1.**
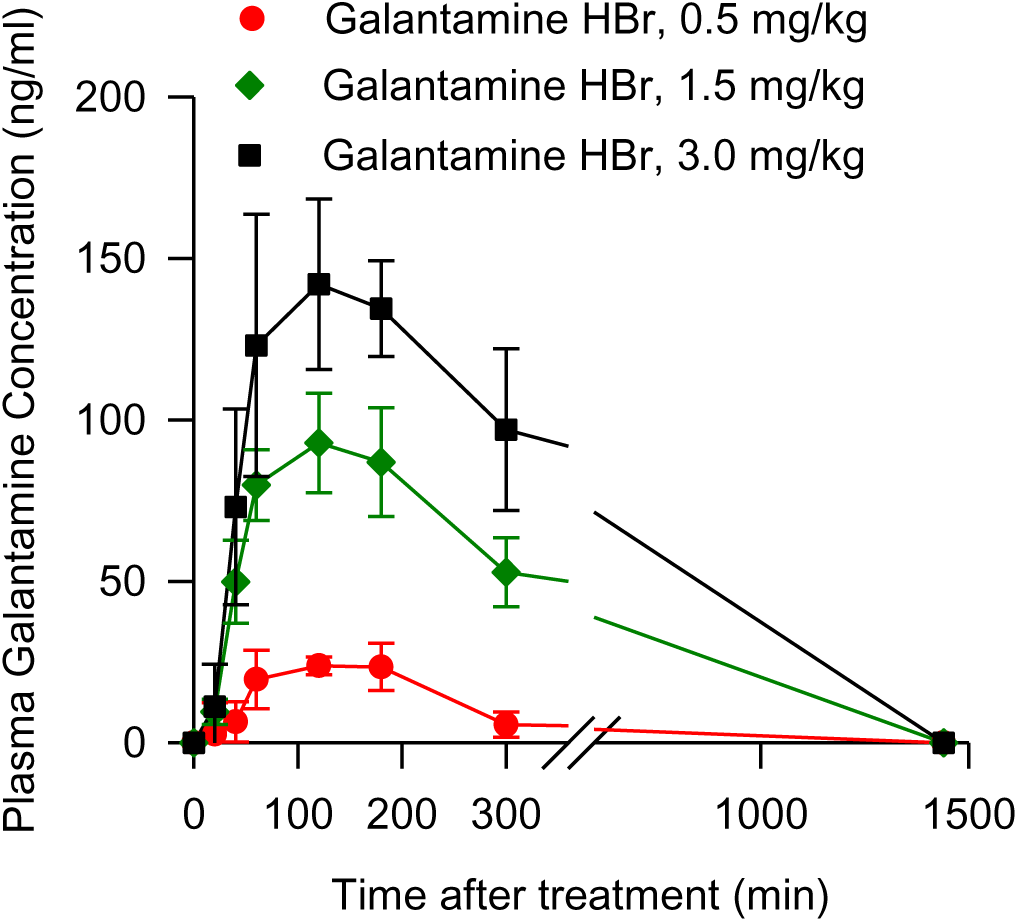
Concentrations of galantamine measured in the plasma obtained at various times from non-human primates treated orally with galantamine. Points and error bars represent mean and SEM, respectively, of results obtained from Cynomolgus monkeys treated with 0.5, 1.5, or 3.0 mg/kg galantamine HBr (n = 5, 5, and 4).

Values of maximal plasma concentration of galantamine (Cmax) and time to reach Cmax (i.e., Tmax) were derived directly from the plasma concentrations measured at the various time points from each animal. Table 1 shows the pharmacokinetic parameters derived for each dose studied here. Statistical analysis of the data using a one-way ANOVA revealed that neither t1/2 nor Tmax varied significantly with the dose of galantamine. However, both Cmax and AUC increased significantly as the dose of galantamine increased from 0.5 to 3.0 mg/kg (Cmax: F(2,13) = 33.19, p < 0.001; AUC: F(2,13) = 11.6, p = 0.002).

**Table 1.** Pharmacokinetic profile of oral galantamine in Cynomolgus monkeys.

|  | Galantamine dose (mg/kg) |  |  |
| --- | --- | --- | --- |
|  | 0.5 mg/kg | 1.5 mg/kg | 3.0 mg/kg |
| C<sub>max</sub> (ng/ml) | 33.6 ± 4.8 | 106.6 ± 14.5** | 186.7 ± 20.3***,†† |
| AUC (ng/ml.h) | 159.7 ± 106.0 | 480.3 ± 24.0* | 701.8 ± 104.2** |
| T<sub>max</sub> (h) | 2.0 ± 0.5 | 1.7 ± 0.6 | 2.8 ± 0.9 |
| t<sub>1/2</sub> (h) | 3.2 ± 2.3 | 3.2 ± 1.7 | 1.7 ± 0.3 |

Results are presented as mean ± SEM of 5, 5, and 4 Cynomolgus monkeys treated orally with 0.5, 1.5, or 3.0 mg/kg galantamine, respectively. One-way ANOVA revealed that C<sub>max</sub> and AUC increased significantly with increasing doses of galantamine (see text). Pairwise comparisons using the Tukey post-hoc test: \*, p < 0.05; \*\*, p < 0.01; \*\*\*, p < 0.001 (comparison with 0.5 mg/kg); ††, p < 0.01 (comparison with 1.5 mg/kg).

As shown in Figures 2A and 2B, a Pearson correlation analysis revealed that increasing doses of galantamine correlated linearly with increasing Cmax (r^2^ = 0.70; p < 0.001) and AUC (r2 = 0.62; p < 0.001). Likewise, as shown in Figure 2C, Cmax and AUC correlated linearly (r^2^ = 0.84; p < 0.001).

**Figure 2.**
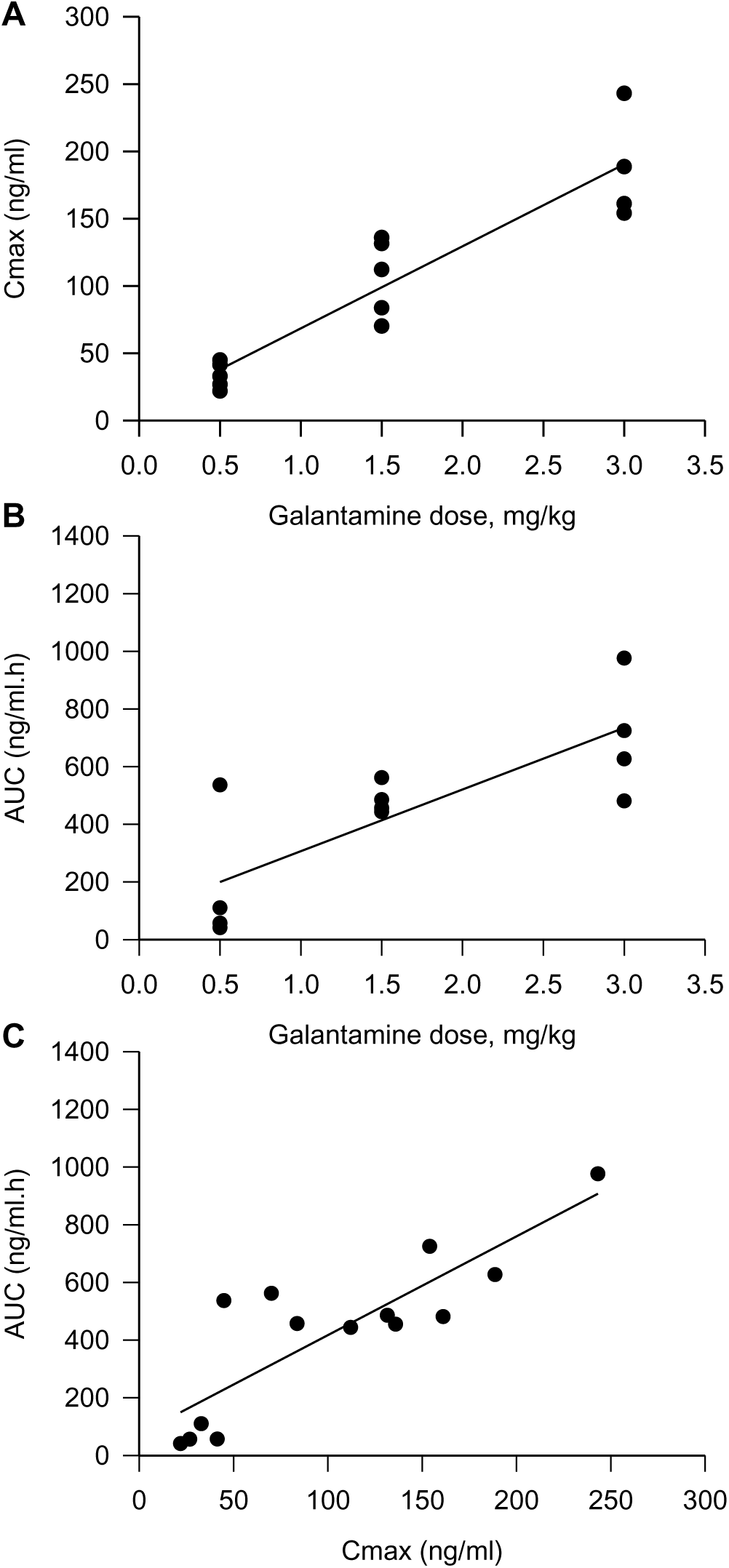
Correlation between doses of galantamine and plasma Cmax or AUC and between Cmax and AUC. ***A.*** Maximal plasma concentrations (Cmax) of galantamine measured after treatment of animals is linearly correlated with the dose of galantamine (r^2^ = 0.70; p < 0.001). ***B.*** Area-under-the curve (AUC) of plasma galantamine concentrations vs. time is linearly correlated with the dose of galantamine (r^2^ = 0.62; p < 0.001). ***C.*** Cmax and AUC are also linearly correlated. In all graphs, each data point represents results obtained from one animal. The solid line represents the linear regression of the data points (r^2^ = 0.84; p < 0.001).

### Blood AChE Activity Following Oral Treatment of Cynomolgus Monkeys with Galantamine HBr

Levels of blood AChE inhibition increased with time after the administration of galantamine HBr to the monkeys, peaked between 1 and 3 h after the treatment, and subsequently declined. At 24 h after the treatment, there was no significant inhibition of blood AChE (Figure 3). The time-dependent rise and fall of blood AChE inhibition paralleled the rise and fall of the plasma galantamine concentration (see Figure 1 and Figure 3), as anticipated based on the reversibility of AChE inhibition by galantamine.

**Figure 3.**
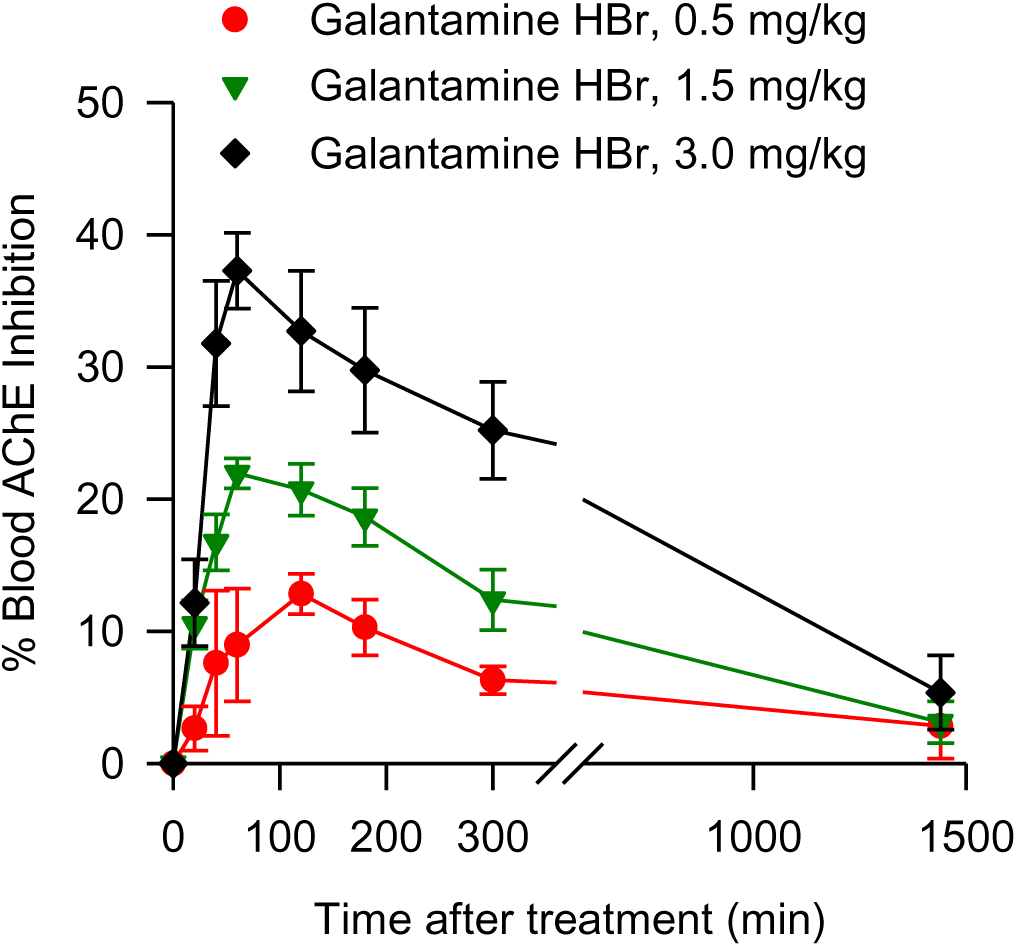
Degree of AChE inhibition measured in whole blood obtained at various times from non-human primates treated orally with galantamine. Points and error bars represent mean and SEM, respectively, of results obtained from the same Cynomolgus monkeys as those in Figure 1.

To determine whether the degree of AChE inhibition measured in the blood of galantamine-treated monkeys was driven by the plasma concentrations of galantamine in these animals, AChE activity was analyzed in whole blood that was drawn from untreated monkeys and spiked *in vitro* with known concentrations of galantamine. As shown in Figure 4A, blood AChE activity decreased as the concentrations of galantamine increased from 0.01 to 10 µM. Fitting of the plot of galantamine concentrations *vs* blood AChE activity with the Hill equation revealed that, under the experimental conditions used in this study, the IC50 for galantamine to inhibit AChE activity in blood of Cynomolgus monkeys is 1.55 ± 0.15 µM. The curve fitting was then used to estimate the degree of AChE inhibition produced by the maximal concentrations of galantamine measured in the plasma of treated animals. The finding that the estimated % AChE inhibition by maximal plasma concentrations of galantamine measured in treated animals significantly correlated with the percentage of AChE inhibition measured in the blood of those animals (r^2^ = 0.56, p = 0.001) confirmed that the degree of AChE inhibition measured in the blood of galantamine-treated animals depends on the blood concentrations of galantamine (Figure 4B).

**Figure 4.**
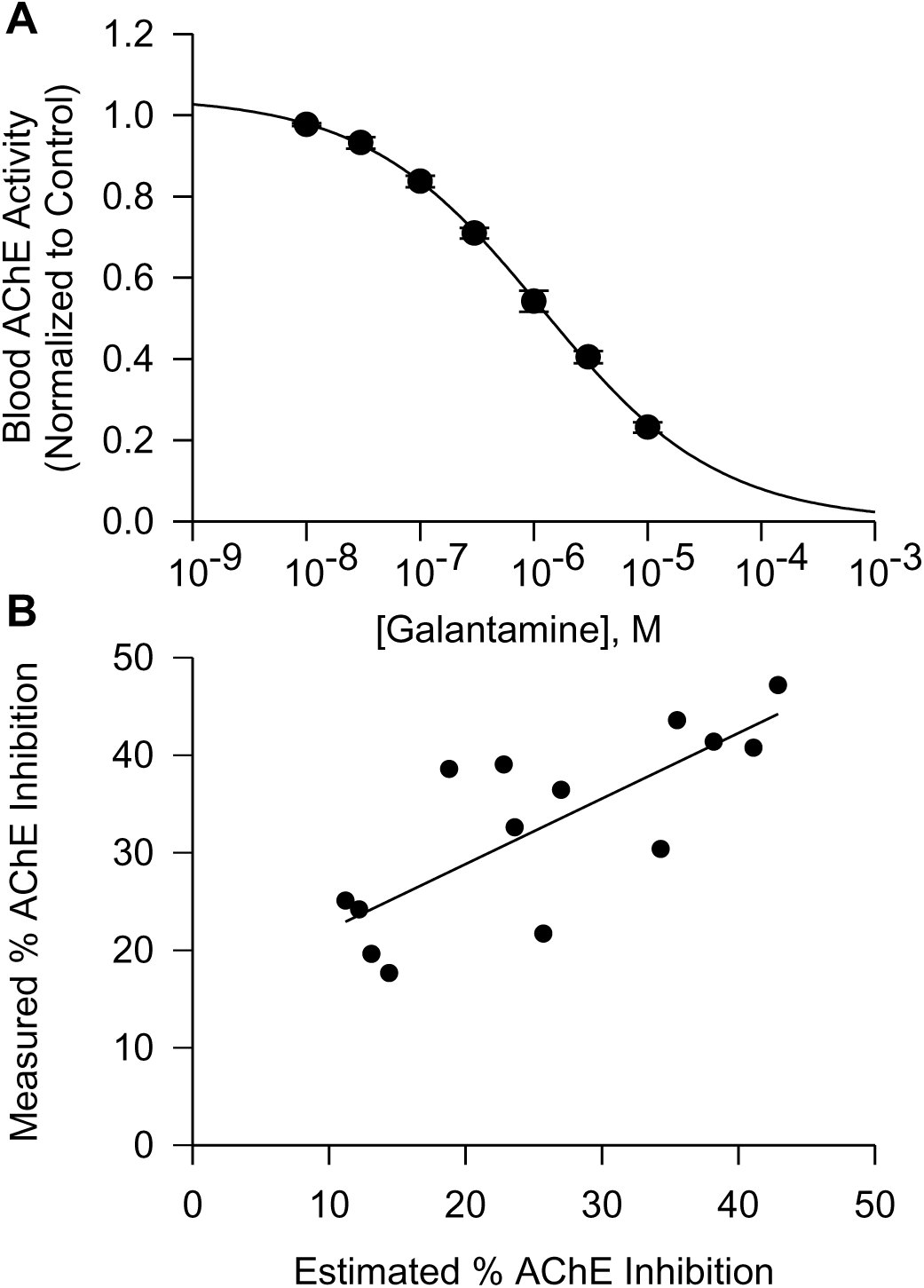
Concentration-dependent inhibition of blood AChE by galantamine in vitro and correlation between the degree of AChE inhibition measured in blood of galantamine-treated animals and the degree of AChE inhibition estimated to be generated by plasma concentrations of galantamine in the treated animals. ***A.*** Concentration-response relationship for galantamine-induced inhibition of AChE in whole blood of Cynomolgus monkeys. Data points and error bars represent mean and SEM, respectively, of results obtained from triplicate assays. ***B.*** Maximal degree of AChE inhibition measured in blood of Cynomolgus monkeys treated with 0.5, 1.5, or 3.0 mg/kg galantamine correlated linearly with the degree of AChE inhibition estimated to be produced by the peak concentrations of galantamine measured in the plasma of the treated animals (r^2^ = 0.56, p = 0.001). Each data point represents results obtained from one animal, and the solid line represents the linear regression of the data points.

### Effectiveness of oral galantamine pretreatment to counter soman-induced acute toxicity in Cynomolgus monkeys post-treated with conventional antidotes

The severity of the intoxication induced by 4.0xLD50 soman in non-human primates post-treated with atropine, 2-PAM, and midazolam was evaluated using a qualitative scoring system based on the clinical neurological signs recorded in the FOB. A toxicity score of 1 was defined by a period of hypoactivity and/or low responsiveness to an external stimulus, with animals presenting full recovery within 1-2 h after the treatments. A score of 2 was defined by clonic tremors or short-lasting (<10 min) mild whole-body motor tonic-clonic convulsions that did not recur within 24 h, with animals presenting full recovery within 3-4 h after the treatments. A score of 3 was defined by clonic tremors or mild tonic-clonic convulsions that also lasted <10 min but recurred 2-3 times within 1-2 h after onset and were accompanied by muscle fasciculations, occasional nystagmus, and loss of posture, with animals showing signs of full recovery within 3-4 h after the treatments. A score of 4 was defined by mixed periods of long lasting (10-20 min) whole-body motor convulsions and hypoactivity throughout the day, muscle fasciculations, and loss of posture, with some animals showing short-lasting periods of a comatose state after bouts of very strong convulsions. Animals could still recover from a toxicity score 4, as evidenced by periods of normal posture and normal responsiveness that were evident starting between 4 and 8 h after the treatments. A score of 5, on the other hand, was defined by life-threatening signs of intoxication from which the animals were unlikely to recover. These signs included unremitting convulsions lasting >20 min, lack of response to external stimuli, severe muscle fasciculations, loss of posture, severe respiratory distress characterized by gasping, and low body temperature by palpation. As soon as animals presented these life-threatening signs of intoxication, they were humanely euthanized according to the IACUC-approved protocol.

All four Cynomolgus monkeys pre-treated orally with saline, injected intramuscularly with 4.0xLD50 soman, and post-treated intramuscularly with atropine, 2-PAM, and midazolam presented clinical signs of intoxication within 10-15 min after the soman challenge. Typical signs of intoxication included rapid onset of hyperexcitation, characterized by initial agitation that quickly evolved to frantic movement and jumps. In all animals, the severity of the intoxication evolved quickly. Specifically, between 15 and 20 min after the soman injection, intermittent clonic tremors developed and progressed to whole-body tonic-clonic motor convulsions. Approximately 20-30 min after the soman challenge, this severe excitatory phase gave way to a short-lasting comatose state that was followed by periods of hyper- and hyporeactivity. Approximately 1 h after the soman challenge, motor convulsions recurred in all saline-pretreated animals. As time progressed, recurring convulsions increased in severity and duration and were accompanied by muscle fasciculations, salivation, lacrimation, increased urination, diarrhea, nystagmus, loss of posture, loss of body temperature, and severe respiratory distress characterized by gasping. Between 90 min and 4 h after the soman challenge, all saline-pretreated animals presented a toxicity score of 5, which defined the endpoint for humane euthanasia (Figure 5A, 5B).

**Figure 5.**
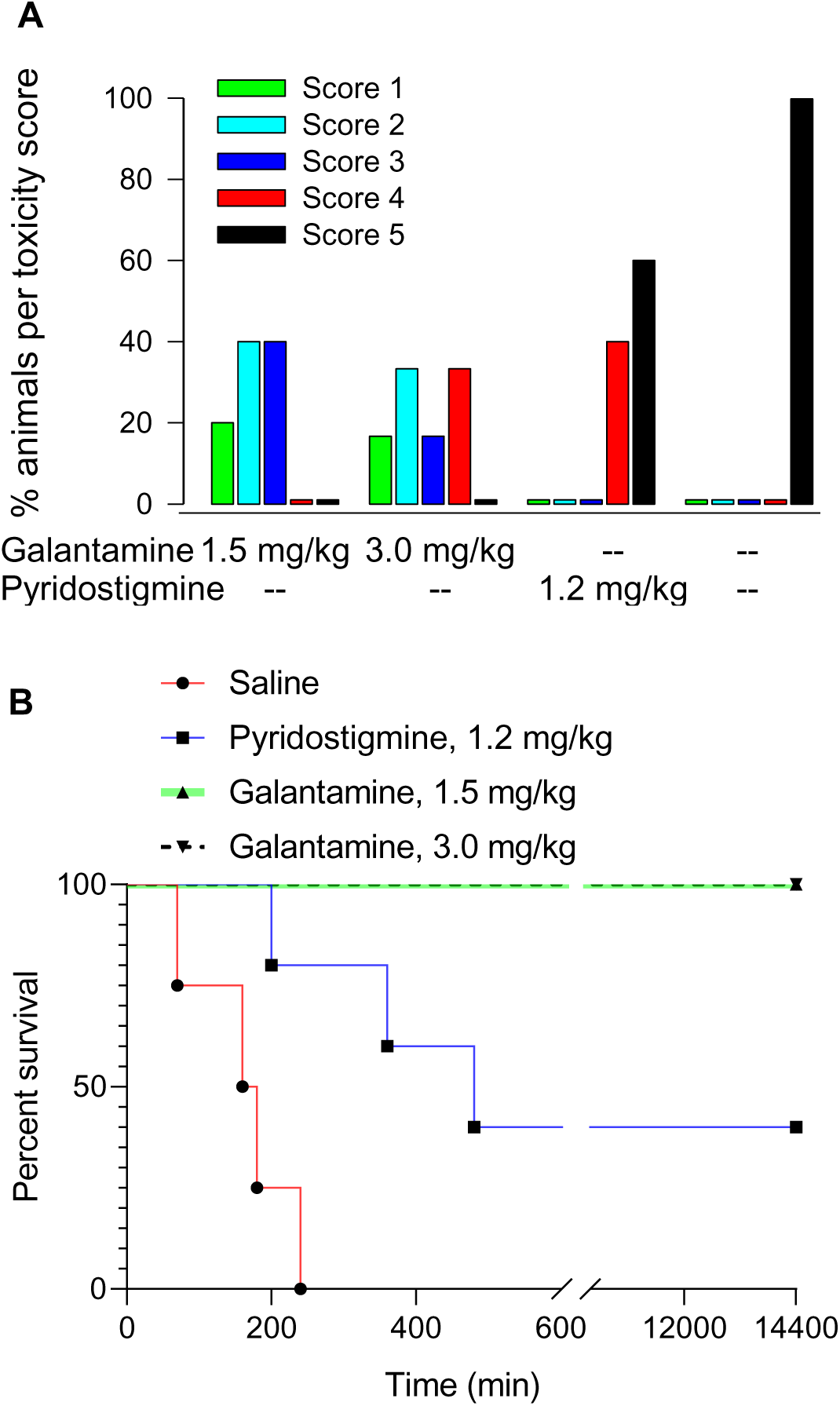
Effectiveness of pretreatment with galantamine, pyridostigmine, or saline to mitigate the acute toxicity of 4.0xLD50 soman in Cynomolgus monkeys post-treated with atropine, 2-PAM, and midazolam. *A.* Percentage of animals presenting a given toxicity score following their challenge with 4.0xLD50 soman when they were pretreated with galantamine HBr (1.5 or 3.0 mg/kg, n = 5 and 6), pyridostigmine (1.2 mg/kg, n = 5), or saline (n = 4) and post-treated with atropine (0.4 mg/kg), 2-PAM (30 mg/kg), and midazolam (0.32 mg/kg). The greater the score is, the more severe the signs of acute toxicity are. *B.* Kaplan-Meier plots of the survival of the soman-challenged animals subjected to the different pre-treatments. The survival curves obtained from the three pretreatment groups were significantly different (χ^2^ = 26.84; df = 3, p < 0.0001).

Pyridostigmine-pretreated monkeys that were injected with 4.0xLD50 soman and post-treated with conventional antidotes also developed typical clinical signs of intoxication within 10-15 min after the soman challenge. However, in two animals, the pyridostigmine pretreatment prevented clinical signs of intoxication from becoming life-threatening. In these two animals, clinicals signs of intoxication progressed from the rapid onset of hyperexcitation to intermittent clonic tremors followed by bouts of whole-body motor convulsions lasting 15-20 min that, as in the saline-pretreated animals, gave way to a short-lasting comatose state. Approximately 1 h after the soman challenge, motor convulsions recurred in these two pyridostigmine-pretreated animals, which also presented muscle fasciculations, salivation, lacrimation, increased urination, and diarrhea. Clinical signs of intoxication did not progress further in these two pyridostigmine-pretreated animals, which survived for 24 h after the soman challenge. Between 3 and 8 h after the soman challenge, the other three pyridostigmine-pretreated monkeys had to be euthanized as they presented the life-threatening signs of intoxication that defined the toxicity score of 5 (Figure 5A, 5B).

Pretreatment with 1.5 or 3.0 mg/kg galantamine delayed by 30-40 min the onset and reduced the severity of clinical signs of toxicity in all soman-challenged monkeys that were post-treated with the conventional antidotes. In the group of five animals that had been pre-treated with 1.5 mg/kg galantamine HBr, challenged with 4.0xLD50 soman, and post-treated with conventional antidotes, one, two, and two animals presented the clinical signs that defined a toxicity score of 1, 2, and 3, respectively (Figure 5A). In the group of six animals that had been pretreated with 3mg/kg galantamine HBr, one, two, one, and two animals presented the clinical signs that defined the toxicity scores of 1, 2, 3, and 4, respectively (Figure 5A). No animal in these groups presented a toxicity score of 5. At approximately 4-6 h after the soman challenge, all animals pretreated with 1.5 or 3.0 mg/kg galantamine were no longer presenting clinical signs of toxicity. All galantamine-pretreated animals survived 24 h after the soman challenge (Figure 5B).

A Kaplan-Meier analysis revealed that the survival curves obtained from the three pretreatment groups were significantly different (χ^2^ = 26.84; df = 3, p < 0.0001) (Figure 5B). Pairwise comparisons revealed that % survival of soman-challenged monkeys that were pretreated with saline and post-treated with conventional antidotes was significantly lower than % survival of animals that had been pretreated with 1.5 mg/kg galantamine (uncorrected p = 0.0027; Bonferroni-corrected p = 0.0135) or 3.0 mg/kg galantamine (uncorrected p = 0.0011; Bonferroni-corrected p = 0.0055). Based on uncorrected p values, % survival of soman- challenged monkeys pretreated with saline was also lower than % survival of animals pretreated with pyridostigmine (p = 0.0108). In addition, % survival of animals pretreated with pyridostigmine was lower than % survival of animals pretreated with 1.5 mg/kg galantamine (p = 0.034) or 3.0 mg/kg galantamine (p = 0.049).

### Effectiveness of oral galantamine pretreatment to counter soman-induced cytotoxicity in the brain of Cynomolgus monkeys post-treated with conventional antidotes

Cytotoxicity was evaluated in brain sections that were stained with FJB, a high-affinity fluorescent marker for localization of neurodegeneration during acute neuronal distress (Schmued and Hopkins, 2000). As shown in Figure 6, large numbers of FJB-positive cells were visualized in the CA1 field of the hippocampus of animals that were treated with saline or pyridostigmine before their exposure to soman followed by immediate post-treatment with atropine, 2-PAM, and midazolam. In contrast, very few FJB-positive cells were detected in the hippocampus of animals that were pretreated with galantamine, challenged with 4.0xLD50 soman, and post-treated with atropine, 2-PAM, and midazolam.

**Figure 6.**
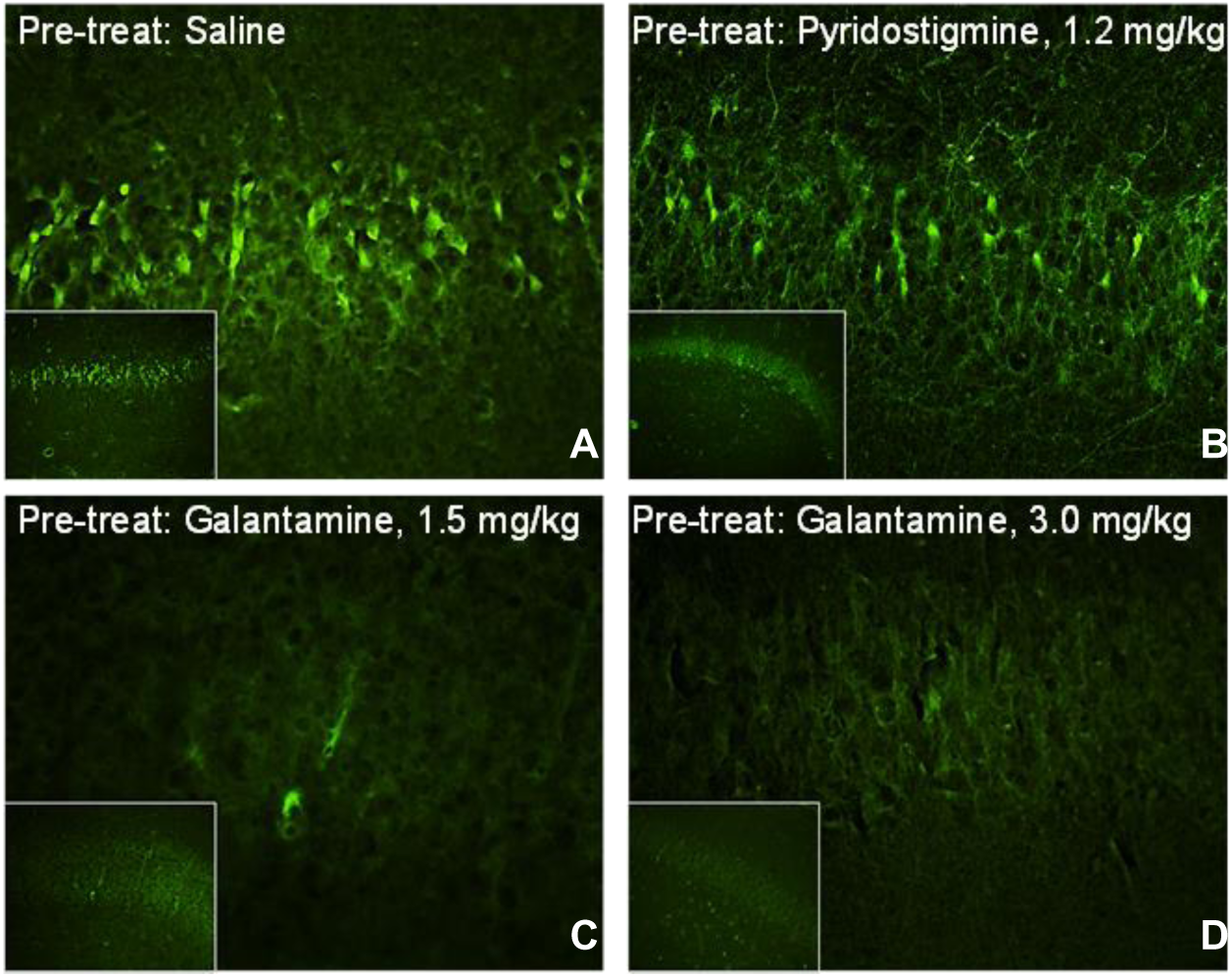
Neurodegeneration in the hippocampi of soman-challenged Cynomolgus that were pretreated with galantamine, pyridostigmine, or saline and post-treated with conventional antidotal therapy. ***Panel A.*** Large numbers of Fluoro Jade-B (FJ-B)-positive (dead) cells were visualized in the hippocampi of Cynomolgus monkeys that were pretreated orally with saline, challenged with 4.0xLD50 soman, and post-treated intramuscularly with 0.4 mg/kg atropine, 30 mg/kg 2-PAM, and 0.32 mg/kg midazolam. Similar results were obtained from Cynomolgus that were pretreated orally with 1.2 mg/kg pyridostigmine bromide and euthanized 24 h after the soman challenge (***panel B***) or soon after presenting life-threatening signs of intoxication. Very few FJB-positive cells were visualized in the hippocampi of galantamine (1.5 or 3.0 mg/kg, oral)-pretreated animals that were euthanized 24 h after their challenge with soman (***panels C and D)***.

## Discussion

The present study is the first to demonstrate that oral pretreatment with galantamine effectively increases the effectiveness of conventional antidotal post-treatment to counter the acute toxicity of 4.0xLD50 soman in Cynomolgus monkeys. Compared to oral pretreatments with saline or pyridostigmine, oral pretreatment with clinically relevant doses of galantamine significantly increased survival of soman-challenged non-human primates post-treated with the conventional antidotal therapy consisting of doses of atropine, 2-PAM, and midazolam equivalent to those recommended for pre-hospital management of nerve agent intoxication. The remarkable ability of galantamine to improve the effectiveness of the conventional antidotal therapy against soman is discussed herein.

In 2003, the FDA approved the use of pyridostigmine, a reversible AChE inhibitor commonly prescribed to treat Myasthenia gravis, as pretreatment for military personal at risk of exposure to the nerve agent soman (Aebersol, 2012). The approval was granted following the demonstration that pretreatment with pyridostigmine significantly increased survival of guinea pigs and non-human primates exposed to supra-lethal doses of the nerve agent soman and post-treated with atropine, oximes, and benzodiazepines (see Lorke and Petroianu, 2019 and references therein). For instance, Rhesus monkeys that were challenged with 5.0xLD50 soman and post-treated with atropine (0.4 mg/kg), 2-PAM (25.71 mg/kg), and diazepam or midazolam (1 mg/kg) survived for 48 h when they were pretreated orally with pyridostigmine bromide (1.2 mg/kg), although they still presented temporary incapacitation and significant neurodegeneration in different brain regions (Hayward et al., 1990). Similar results were obtained in the present study, which used a lower dose of midazolam to mimic the doses recommended for pre-hospital management of nerve agent poisoning. Specifically, following oral pretreatment with 1.2 mg/kg pyridostigmine bromide, 2 out of 5 soman (4.0xLD50)-challenged Cynomolgus monkeys post-treated with atropine (0.4 mg/kg), 2-PAM (30 mg/kg), and midazolam (0.32 mg/kg) survived for 24 h post-challenge. By contrast, none of the Cynomolgus monkeys survived when they were pretreated with saline instead of pyridostigmine, challenged with 4.0xLD50 soman, and post-treated with the conventional antidotes. However, regardless of their pretreatment with pyridostigmine, neurodegeneration was still detected in the hippocampi of Cynomolgus monkeys that survived for 24 h after the soman challenge.

The effectiveness of pyridostigmine as a pretreatment against nerve agent poisoning has been attributed to its ability to reversibly inhibit AChE and, consequently, protect a critical portion of the enzyme from irreversible inhibition by soman (Dinhurber et al., 1979; Gordon et al., 1978; Haigh et al., 2005; Heyl et al., 1980; Eckert et al., 2006). However, because pyridostigmine does not promptly cross the blood brain barrier, it does not protect AChE in the brain from irreversible inhibition by soman and, consequently, it does not prevent the CNS-related toxic effects of this agent (Anderson et al., 1992). Although centrally acting, reversible AChE inhibitors, including galantamine and huperzine, are more effective pretreatments than pyridostigmine to prevent the acute toxicity of soman and other nerve agents (Albuquerque et al., 2006; Hilmas et al., 2009; Hamilton et al., 2017), concerns have been raised that effective doses of centrally acting reversible AChE inhibitors can be detrimental to executive brain functions.

In the present study, all Cynomolgus monkeys orally pretreated with galantamine (1.5 or 3.0 mg/kg) survived a subsequent challenge with 4.0xLD50 soman when post-treated with conventional antidotal therapy consisting of atropine (0.4 mg/kg, i.m.), 2-PAM (30 mg/kg, i.m.), and midazolam (0.32 mg/kg). In addition, no neurodegeneration was detected in the hippocampi of the galantamine-pretreated Cynomolgus monkeys. The effective doses of galantamine were found to reversibly inhibit blood AChE activity by approximately 20%-40%, a degree of inhibition that: (i) is observed in humans treated with clinically recommended doses of galantamine (Bickel et al., 1991), and (ii) is reported to be safe and sufficient to prevent the acute toxicity of soman (Dunn and Sidell, 1989). The higher effectiveness of galantamine compared to that of pyridostigmine may be accounted for by its ability to cross the blood-brain-barrier and protect not only peripheral, but also central AChE from the irreversible inhibition by soman. However, the possibility cannot rule out that additional actions account for the higher effectiveness of galantamine compared to that of pyridostigmine. For instance, in contrast to pyridostigmine, galantamine has higher potency to inhibit AChE than butyrylcholinesterase, a natural OP scavenger. In addition, AChE-unrelated neuroprotective properties of galantamine may be important determinants of its antidotal effectiveness against nerve agents (reviewed in Pereira et al., 2010).

In a previous study, a single intramuscular dose of 0.362 mg/kg galantamine, which was estimated to produce approximately 50% inhibition of blood AChE activity, also protected Cynomolgus monkeys against the acute toxicity of 1.0xLD50 soman in the absence of any antidotal posttreatment (Hamilton et al., 2017). In that study, too, galantamine was found to be superior to pyridostigmine as a pretreatment against soman intoxication. As importantly, however, Hamilton et al. (2017) also reported that the effective dose of galantamine to protect Cynomolgus monkeys against the toxicity of 1.0xLD50 soman had no detrimental effect on the cognitive function of the non-human primates. Only doses that were ≥40% higher than the effective dose induced a transient, yet statistically significant cognitive deficit in Cynomolgus monkeys (Hamilton et al., 2017).

Use of pharmacokinetic parameters and pharmacodynamics measures provides a well-accepted method for conversion of doses from one animal species to another (Reigner and Blesch, 2002). This was the approach used by the FDA to convert oral doses of pyridostigmine HBr that prevented the acute toxicity of soman in non-human primate to human equivalent doses (HEDs) to be used by military personnel at risk of exposure to soman. Here, effective doses of galantamine HBr to prevent soman lethality were translated to human equivalent doses using the pharmacokinetic profile of oral galantamine in these animals and in humans.

A single oral treatment of humans with 8 mg galantamine produces an AUC of 406 ± 77 ng*h/ml (Huang, 2010; Zhao et al., 2002). By comparison, a single oral treatment of Cynomolgus monkeys with 1.5 or 3.0 mg/kg galantamine HBr, which effectively protected the animals against the acute toxicity of 4.0xLD50 soman, produced AUCs of 480 ± 24 ng*h/ml and 702 ± 104 ng*h/ml, respectively (Table 1). Using the equation *HED = AUC_animalx_Cl_human_*, where AUC_animal_ is the AUC produced by the target dose in an animal and CL_human_ is the clearance of galantamine in humans, it was possible to estimate the HED of galantamine needed to prevent the acute toxicity of soman in humans. Considering that CL of galantamine in humans is approximately 19 l/h, the oral doses of 1.5 and 3.0 mg/kg galantamine HBr in Cynomolgus monkeys translate to approximately 9.4 and 13.7 mg galantamine base in humans. These doses are clinically relevant because daily oral doses of galantamine base in the range of 8 mg to 24 mg are currently approved for treatment of Alzheimer’s disease, and, therefore, considerable post-market safety data is available for these doses. In addition, a recent clinical study reported no clinical signs of toxicity and no detrimental cognitive effect in healthy human subjects treated with a single dose of 8 mg galantamine (Morasch et al., 2015).

In conclusion, the data presented here demonstrate that non-human primates are effectively protected against the acute toxicity of soman when they are pretreated with a clinically relevant oral dose of galantamine and post-treatment with doses of conventional antidotes recommended for pre-hospital management of nerve agent poisoning. They also support the contention that this antidotal therapy is more effective than the standard post-treatments alone or in association with pyridostigmine as pretreatment. Therefore, this study lays the groundwork for the continued advanced development of galantamine as a safe and effective oral pretreatment against OP poisoning.

## Acknowledgments

The authors are indebted to Ms. Mabel A. Zelle for her technical assistance. This work was supported in part by the Countervail Corporation with funds from the CounterACT Program of the National Institutes of Neurological Disorders and Stroke via a Phase II SBIR grant [R44 NS068049].

## Conflict of Interest Statement

The use of galantamine as an antidote against OP poisoning is protected under the Patent Application PCT/US05/33789 filed by the University of Maryland Baltimore and the U.S. Medical Research Institute of Chemical Defense on 2005 Sept 23 and licensed to Countervail Corp.

## References

Aebersold P. FDA experience with medical countermeasures under the animal rule. Adv Prev Med 2012:507571, 2012.

Albuquerque EX, Pereira EFR, Aracava Y, Fawcett WP, Oliveira M, Randall WR, Hamilton TA, Kan RK, Romano JA Jr, Adler M. Effective countermeasure against poisoning by organophosphorus insecticides and nerve agents. Proc Natl Acad Sci U S A 103:13220–13225, 2006.

Anderson DR, Harris LW, Woodard CL, Lennox WJ. The effect of pyridostigmine pretreatment on oxime efficacy against intoxication by soman or VX in rats. Drug Chem Toxicol 15:285–94, 1992.

Bickel U, Thomsen T, Weber W, Fischer JP, Bachus R, Nitz M, Kewitz H. Pharmacokinetics of galanthamine in humans and corresponding cholinesterase inhibition. Clin Pharmacol Ther 50: 420–428, 1991.

Chai PR, Boyer EW, Al-Nahhas H, Erickson TB. Toxic chemical weapons of assassination and warfare: nerve agents VX and sarin. Toxicol Commun 1:21–23, 2017.

Dirnhuber P, French MC, Green DM, Leadbeater L, Stratton JA. The protection of primates against soman poisoning by pretreatment with pyridostigmine. J Pharm Pharmacol 31:295–299, 1979.

Dunn MA, Sidell FR. Progress in medical defense against nerve agents. JAMA. 262:649–652, 1989.

Eckert S, Eyer P, Mückter H, Worek F. Kinetic analysis of the protection afforded by reversible inhibitors against irreversible inhibition of acetylcholinesterase by highly toxic organophosphorus compounds. Biochem Pharmacol 72:344–357, 2006.

Franca TCC, Kitagawa DAS, Cavalcante SFA, da Silva JAV, Nepovimova E, Kuca K. Novichoks: The dangerous fourth generation of chemical weapons. Int J Mol Sci 20(5). pii: E1222, 2019.

Gauvin DV, Baird TJ. A functional observational battery in non-human primates for regulatory-required neurobehavioral assessments. J Pharmacol Toxicol Methods 58:88–93, 2008.

Golime R, Palit M, Acharya J, Dubey DK. Neuroprotective effects of galantamine on nerve agent-induced neuroglial and biochemical changes. Neurotox Res 33:738–748, 2018.

Gordon JJ, Leadbeater L, Maidment MP. The protection of animals against organophosphate poisoning by pretreatment with a carbamate. Toxicol Appl Pharmacol 43:207–216, 1978.

Haigh JR, Johnston SR, Peters BM, Doctor BP, Gordon RK, Adler M, Gall KJ, Deshpande SS. Inhibition of guinea pig hemi-diaphragm acetylcholinesterase activity by pyridostigmine bromide and protection against soman toxicity. Chem Biol Interact 157-158:381–382, 2005.

Hamilton LR, Schachter SC, Myers TM. Time course, behavioral safety, and protective efficacy of centrally active reversible acetylcholinesterase inhibitors in Cynomolgus macaques. Neurochem Res 42:1962–1971, 2017.

Hayward IJ, Wall HG, Jaax NK, Wade JV, Marlow DD, Nold JB. Decreased brain pathology in organophosphate-exposed rhesus monkeys following benzodiazepine therapy. J Neurol Sci 98:99–106, 1990.

Heyl WC, Harris LW, Stitcher DL. Effects of carbamates on whole blood cholinesterase activity: chemical protection against soman. Drug Chem Toxicol 3:319–332, 1980.

Hilmas CJ, Poole MJ, Finneran K, Clark MG, Williams PT. Galantamine is a novel post-exposure therapeutic against lethal VX challenge. Toxicol Appl Pharmacol 240:166–173, 2009.

Huang F, Fu Y. A review of clinical pharmacokinetics and pharmacodynamics of galantamine, a reversible acetylcholinesterase inhibitor for the treatment of Alzheimer’s disease, in healthy subjects and patients. Curr Clin Pharmacol. 5:115–124, 2010.

Hurst CG, Newmark J, Romano JA., Jr (2012) Chemical terrorism: Introduction, in Harrison’s Principles of Internal Medicine, 18th ed (Longo DL, Fauci AS, Kasper DL, Hauser SL, Jameson JL, Loscalzo J, editors. eds) pp 1779–1788, McGraw-Hill, New York

Jett DA, Laney JW. Civilian research on chemical medical countermeasures. Toxicol Mech Methods Oct2:1–2, 2019.

Johnson CD, Russell RL. A rapid, simple radiometric assay for cholinesterase, suitable for multiple determinations. Anal Biochem 64:229–238, 1975.

Lorke DE, Petroianu GA. Reversible cholinesterase inhibitors as pretreatment for exposure to organophosphates. A review. J Appl Toxicol 39:101–116, 2019.

Mamczarz J, Kulkarni GS, Pereira EF, Albuquerque EX. Galantamine counteracts development of learning impairment in guinea pigs exposed to the organophosphorus poison soman: clinical significance. Neurotoxicology 32:785–798, 2011.

Marrs TC, Rice P, Vale JA. The role of oximes in the treatment of nerve agent poisoning in civilian casualties. Toxicol Rev 25:297–323, 2006.

McDonough JH, McMonagle JD, Shih TM. Time-dependent reduction in the anticonvulsant effectiveness of diazepam against soman-induced seizures in guinea pigs. Drug Chem Toxicol 33:279–283, 2010.

Morasch KC, Aaron CL, Moon JE, Gordon RK. Physiological and neurobehavioral effects of cholinesterase inhibition in healthy adults. Physiol Behav 138:165–172, 2015.

Newmark J. Therapy for acute nerve agent poisoning: An update. Neurol Clin Pract 9:337–342, 2019.

Nishiwaki Y, Maekawa K, Ogawa Y, Asukai N, Minami M, Omae K, Sarin Health Effects Study Group. Effects of sarin on the nervous system in rescue team staff members and police officers 3 years after the Tokyo subway sarin attack. Environ Health Perspect 109:1169–1173, 2001.

Pereira EF, Aracava Y, Alkondon M, Akkerman M, Merchenthaler I, Albuquerque EX. Molecular and cellular actions of galantamine: clinical implications for treatment of organophosphorus poisoning. J Mol Neurosci 40:196–203, 2010.

Reagan-Shaw S, Nihal M, Ahmad N. Dose translation from animal to human studies revisited. FASEB J. 22:659–661, 2008.

Reigner BG, Blesch KS. Estimating the starting dose for entry into humans: principles and practice. Eur J Clin Pharmacol. 57:835–845, 2002.

Romano JA, Jr, King JM. Psychological casualties resulting from chemical and biological weapons. Mil Med 166:21–22, 2001.

Schmued LC, Hopkins KJ. Fluoro-Jade B: a high affinity fluorescent marker for the localization of neuronal degeneration. Brain Res 874:123–130, 2000.

Steiner WE, Pikalov IA, Williams PT, English WA, Hilmas CJ. An extraction assay analysis for galanthamine in guinea pig plasma and its Application to nerve agent countermeasures. J Anal Bioanal Techniques 3:5, 2012.

Suresh PS, Mullangi R, Sukumaran SK. Highly sensitive LC-MS/MS method for determination of galantamine in rat plasma: application to pharmacokinetic studies in rats. Biomed Chromatogr 28:1633–1640, 2014.

Zhao Q, Brett M, Van Osselaer N, Huang F, Raoult A, Van Peer A, Verhaeghe T, Hust R. Galantamine pharmacokinetics, safety, and tolerability profiles are similar in healthy Caucasian and Japanese subjects. J Clin Pharmacol 42:1002–1010, 2002.

